# Recombination and incomplete lineage sorting resolve the enigma of lysozyme evolution

**DOI:** 10.64898/2026.05.31.729045

**Authors:** Dawson Houghtaling, Edward L. Braun, Rebecca T. Kimball

## Abstract

Lysozyme has long been a model for understanding enzyme structure, function, and evolution. Early studies revealed a conflict between organismal phylogeny and the distribution of three functionally important amino acid residues in galliform birds. Lysozymes differing at all three amino acids appear functionally equivalent, but intermediates exhibit reduced stability and have not been observed in nature. However, the phylogeny suggests two independent occurrences of the three mutations, requiring two separate transitions through low fitness intermediates. We reexamined this apparent paradox using phylogenomic methods, accounting for incomplete lineage sorting and intralocus recombination. The lysozyme locus tree conflicts with an estimated species tree, but the conflict involves a short branch in coalescent units, consistent with incomplete lineage sorting. We also found evidence for recombination, with different parts of the lysozyme locus supporting alternative relationships. The three amino acids are encoded by exons located in different recombination-defined segments with different evolutionary histories. These results support a model where ancestral polymorphism, coupled with recombination between independently arising mutations, allowed rapid transitions between peaks in the fitness landscape without fixation of deleterious intermediates. Our findings resolve this long-standing question in lysozyme evolution and highlight the importance of considering complex genealogical processes such as incomplete lineage sorting and recombination when reconstructing ancestral proteins and interpreting apparent cases of molecular convergence.

## Introduction

Since its discovery by Alexander Fleming in 1922 (Fleming 1922), lysozyme has featured prominently in early discoveries about enzymes. It was among the first enzyme with a complete amino acid sequence (Canfield 1963), the first enzyme whose three-dimensional structure was solved (Blake et al. 1965), and the first for which a detailed mechanism of action was proposed (Phillips 1966; Blake et al. 1967). Thus, it was also an enzyme for which comparative data from a variety of species was obtained early on (Jollès et al. 1976; Ibrahimi et al. 1979), allowing exploration of its evolution. Examination of these early sequences highlighted a conflict between the assumed organismal phylogeny and the distribution of key changes among taxa (Jollès et al. 1976; Ibrahimi et al. 1979). To better understand this conflict, lysozyme also became among the first proteins for which “resurrected” ancestral proteins were subjected to biochemical analyses (Malcolm et al. 1990) to better understand its evolutionary history.

This observed conflict between organismal phylogeny and the apparent history of the enzyme involved the avian order Galliformes, which includes well-studied and economically important taxa such as chickens, turkeys, quail, and guineafowl. While overall the amino acid sequences of lysozymes in galliforms exhibit little variation among species, there are three residues buried in the core of the protein structure (Fig. 1a) that vary consistently among clades. The amino acids in this variable triplet are encoded by two different exons (Fig. 1b), but are adjacent in the three-dimensional structure. The two different variants are: 1) TIS (Thr 40, Ile 55, and Ser 91) and 2) SVT (Ser 40, Val 55, or Thr 91). The TIS variant is present in the two deep-branching galliform families, Megapodiidae (megapodes), and Cracidae (guans and chachalacas) and the species-rich Phasianidae (pheasants) whereas the SVT variant is present in Odontophoridae (New World quail) and Numididae (guineafowl) (Fig. 1c). Although evolutionary relationships within galliforms has varied, neither older classifications (Peters 1934; Wetmore 1960; Johnsgard 1986) or molecular phylogenies (Kornegay et al. 1993; Cox et al. 2007; Bonilla et al. 2010; Wang et al. 2013; Kimball and Braun 2014; Hosner et al. 2016, 2017; Stiller et al. 2024) have suggested the two clades with the SVT variant are sister taxa.

**Fig. 1.**
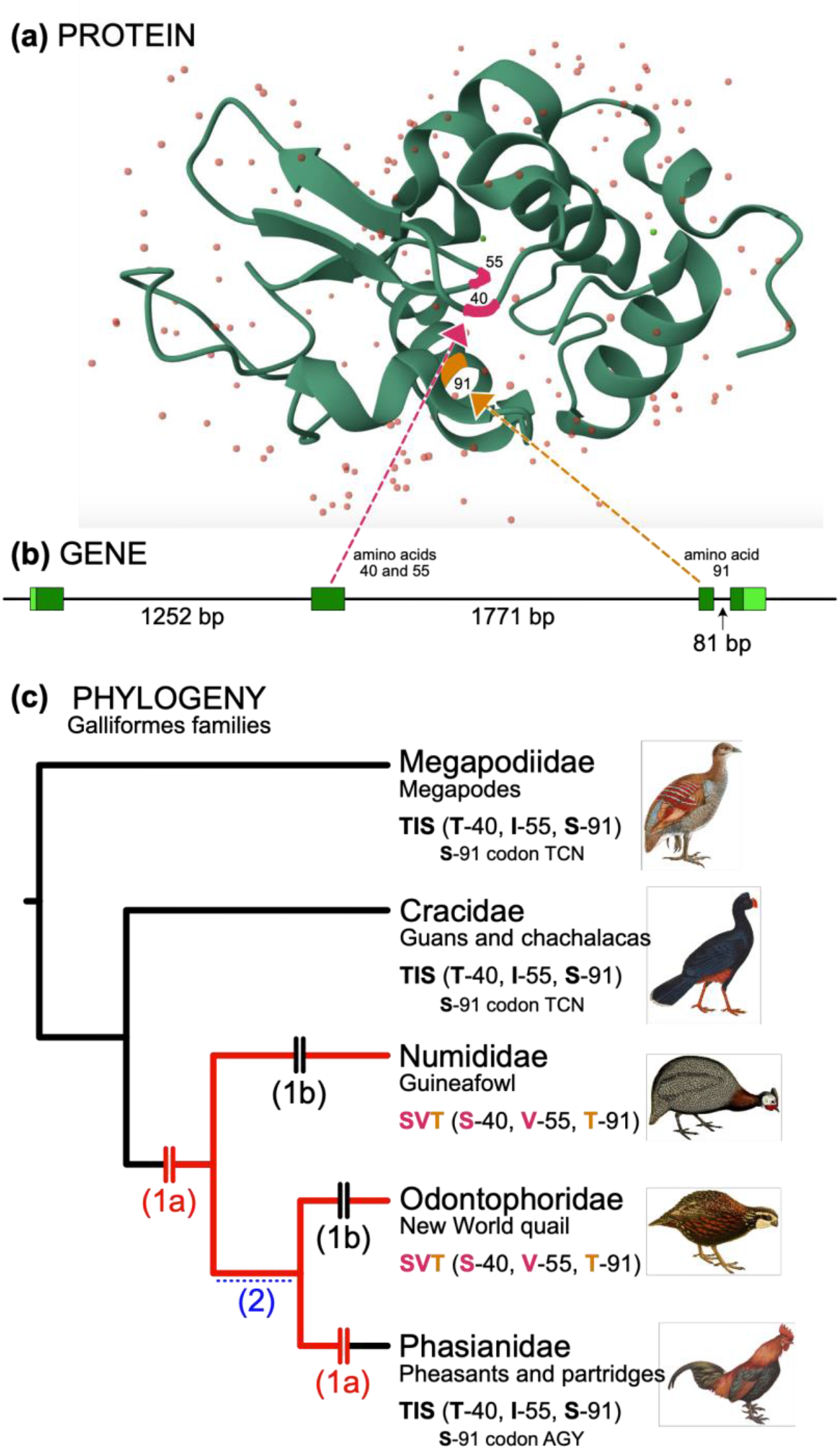
Evolution of the galliform lysozyme c protein. (a) Protein, showing positions of the variable amino acid sites (treating the first residue of the processed protein as residue 1). (b) Intron and exon structure of the galliform *LYZ* gene. The intron sizes are from *Gallus gallus* (chicken); other galliform *LYZ* genes have similar intron sizes. (c) Species tree for the five families of Galliformes. Black branches indicate families with the ancestral state (TIS) and red branches indicate families with the derived state (SVT). The two scenarios for the evolution of *LYZ* if the history of that region matches the species tree are indicated using hash marks on the relevant branches, each with parenthetical numbers. Scenario 1a (red) involves an early TIS→SVT change followed by a reversion to the ancestral state in Phasianidae; scenario 1b (black) involves independent TIS→SVT changes in Numididae and Odontophoridae. The figure is presented as favoring scenario 1a because it is more parsimonious at the level of the codon sequences (the S-91 codon is indicated for families with TIS sequence). Scenario 2 (blue) involves a branch that conflicts between the species tree and the *LYZ* gene tree, with the origin of the SVT allele in the ancestor of Numididae, Odontophoridae, and Phasianidae and retention of a polymorphic state (TIS/SVT) over the branch indicated by the dashed blue line. Scenario 2 requires the serine in TIS to change between a TCN serine codon to a AGY serine codon. Sources of bird images are available in Supplementary File 1

Thus, it is likely that there were at least two evolutionary transitions between the TIS and the SVT variants: either two independent TIS→SVT changes (Scenario 1a in Fig. 1c) or a single TIS→SVT change followed by a reversion of SVT→TIS (Scenario 1b in Fig. 1c). There is indirect evidence that suggests the second scenario is more likely; Kornegay et al. (1993) reported that the serine of a species of chachalaca, from one of the early branching TIS lineages (Cracidae), is encoded by a TCT codon but the serine in the chicken, which is in the derived TIS clade (Phasianidae), is encoded by an AGC codon. This favors the second scenario (a single TIS→SVT followed by a reversal of SVT→TIS) because it involves single nucleotide substitutions whereas the alternative (two TIS→SVT changes) would require either a doublet substitution from a TCN serine codon to an AGY serine codon or two non-synonymous substitutions (either Ser[TCN]→Thr→Ser[AGY] or Ser[TCN]→Cys→Ser[AGY]). Although doublet mutations and indirect mutational pathways are possible, they are rare events (Averof et al. 2000; Rogozin et al. 2016).

Multiple transitions between variants might be likely for proteins that are subject to limited purifying selection or intermediate variants that maintain similar levels of functionality. From its original discovery, lysozyme was known to have antibacterial activity (Fleming 1922; Fleming and Allison 1924), and lysozymes are found throughout the animal kingdom (reviewed by Callewaert and Michiels 2010). In addition to their key role in the innate immune system, lysozyme also appears to be important in facilitating aspects of the acquired immune system (reviewed by Ragland and Criss 2017). Underlying its overall importance, in birds, lysozyme is found in many different tissues throughout the body (Sato and Watanabe 1976; Nile et al. 2004; Vanderven et al. 2012; Wang et al. 2016; Bar Shira and Friedman 2018). It has also long been known to be present in avian egg white (Fleming and Allison 1924), and has been found in eggs from a variety of wild birds (Saino et al. 2002; Wellman-Labadie et al. 2008; D’Alba et al. 2010; Cao et al. 2015) as well as in semen (Sotirov et al. 2002; Rowe et al. 2013) and preen oil (Carneiro et al. 2020). The ubiquity of lysozyme, across the animal kingdom and across many tissues within a species, highlights its importance, and suggests it is likely to be under moderate to strong purifying selection.

In the early comparisons among taxa, no intermediate variants were observed (Jollès et al. 1976; Ibrahimi et al. 1979), suggesting that intermediates likely have lower fitness. To test this, Malcolm et al. (1990) used site-directed mutants to synthesize enzymes with all possible intermediate sequences and examined several metrics of enzyme stability of these different variants. Malcolm et al. (1990) found that the TIS and SVT variants of lysozyme have virtually identical thermal denaturation properties (*T*_m_ = 73.9°C for TIS and *T*_m_ = 73.4°C for SVT; the range between these two was defined as the “neutral zone”) but the range of *T*_m_ values for the single and double mutants that comprise the potential evolutionary steps between those TIS and SVT is much wider (from 70.6°C to 77.5°C) (see Supplementary Fig. S1). Moreover, all pathways between the wild types include at least one intermediate with a *T*_m_ that lies a substantial distance from the wild-type thermostability of 73-74°C. Similar patterns were observed for other metrics considered (Supplementary Fig. S1; Malcolm et al. 1990). If the neutral zone for thermal denaturation defines the range over which enzyme function is favored, then Malcolm et al. (1990) identified a case where enzyme evolution required multiple non-synonymous substitutions with intermediates that are likely to be of lower fitness – twice.

In evolutionary studies, there have been many observations of seemingly complex traits whose distribution among species appears to conflict with known species trees. This is often attributed to independent gains in separate species evolving convergently, potentially reflecting the impact of similar selective pressures that have resulted in similar adaptations. The existence of morphological convergence has been appreciated since the dawn of evolutionary biology as a field (Darwin 1859) and, perhaps unsurprisingly, there are also many examples of convergence at the molecular level (Storz 2016). While Ibrahim et al. (1979) suggested that convergence (or parallelism) might explain the distribution of lysozyme sequences in galliform birds, the possibility that three non-synonymous substitutions converged in two separate exons seems highly unlikely, particularly given that the Malcolm et al. (1990) results suggested intermediates would have been selected against.

Conflicts between gene trees and the species tree are an alternative source of apparent homoplasy known to have an impact both at the level of molecules and of morphological traits (Zelenkov 2011; Wu et al. 2018; Guerrero and Hahn 2018). Discordance between gene trees and the species tree reflects several different processes, including gene duplication and loss, horizontal transfer (i.e., introgressive hybridization), and incomplete lineage sorting (ILS) (Maddison 1997). The last of these three processes is likely to be especially important, since it occurs with a non-zero probability whenever branching evolution occurs with finite population sizes. ILS requires the persistence of ancestral polymorphisms through two or more speciation events (Maddison 1997). Applied to this situation, it would suggest the allele ancestral to the SVT variants arose prior to the divergence of Numididae and Odontophoridae, and was retained as a polymorphism along with the TIS allele ancestral to the lysozyme present in Phasianidae.

Over time, only the SVT allele was fixed in the Numididae and Odontophoridae lineages and the TIS allele was fixed in the ancestor of Phasianidae. This may seem to be a simple explanation for the distribution of the different *LYZ* variants, as it would only require a single transition between the two types. Purifying selection is expected to eliminate deleterious alleles, so loci that are selected against are unlikely to persist in populations long enough to exhibit discordance due to ILS. Thus, it appears that the *LYZ* gene has either transitioned between the variants twice (if its gene tree matches the species tree) or it transitioned once very rapidly and persisted in a polymorphic state through the speciation events leading to Numididae and Odontophoridae (if its gene tree conflicts with the species tree due to ILS).

Here we use modern phylogenomic methods to reexamine the questions raised by the classic Malcolm et al. (1990) study. If the topology of the *LYZ* gene tree matches the species tree we would have to conclude two separate transitions between TIS and SVT (Fig. 1c) led to the current distribution of *LYZ* alleles in galliforms, although the biochemical properties of the reconstructed ancestral genes make this unlikely. The obvious alternative is to postulate that the evolutionary history of all or part of the *LYZ* gene has a different history from that of the species in Galliformes (that ILS explains the current distribution of the TIS and SVT variants). We explored this issue in several ways. First, we estimated the Galliformes species tree using a phylogenomic dataset; this yielded estimates of the coalescent branch lengths (which reflect the probability of discordance due to ILS) and provided information about the fit of the data to a model of evolution by ILS alone (which provides insights into the potential for discordance due to introgression). Second, we examined the complete *LYZ* gene region to determine whether different segments of the *LYZ* gene have distinct histories (i.e., whether the *LYZ* region includes multiple *c*-genes, which correspond to regions with a single gene tree). We complemented this with an examination of whether sites in the *LYZ* locus supported alternative topologies (i.e., whether convergence might lead to sites in the focal amino acids supporting a different topology than the remainder of the locus). We also estimated recombination breakpoints within *LYZ*, and whether recombination might have been important in the TIS to SVT changes. Finally, we discuss the implications of our results for studies that seek to reconstruct ancestral proteins.

## Methods

### Taxon sampling

We included data from 16 galliform species (AviList Core Team 2025) where complete, assembled genomes were available. This included at least one sampled species from each of the five recognized galliform families (Supplementary Table S1). The root of Galliformes is well-established (Hackett et al. 2008; Prum et al. 2015; Kimball et al. 2019; Braun et al. 2024; Stiller et al. 2024) so we did not include an non-galliform outgroup.

### Estimation of a galliform species tree

To estimate a species tree using data independent of the *LYZ* locus, we extracted ultra-conserved element sequences (UCEs; Faircloth et al. 2012) from the published genomes that we used. The UCEs were extracted using PHYLUCE (Faircloth 2016), using the 5K probe set (Sun et al. 2014; available at https://www.ultraconserved.org/). We extracted 1000 bases of flanking sequences on each side of the conserved UCE core. We aligned sequences using MAFFT 7.505 (Katoh and Standley 2013), and trimmed alignments to sites present in at least 50% of taxa. Gene trees were estimated using IQ-TREE 2.1.3 (Minh et al. 2020) with the “TEST” option (Kalyaanamoorthy et al. 2017) to identify the best-fitting model of those available in jModelTest (Posada 2008). Support was estimated using the aLRT (Anisimova and Gascuel 2006). The species tree was estimated using weighted ASTRAL (wASTRAL) version 1.7.2.3 (Zhang and Mirarab 2022) using the hybrid model (incorporating both branch lengths and support).

### LYZ sequence data and alignment

We obtained the complete *LYZ* gene region (exons, introns, and 2000 bp upstream of the start codon) from *Gallus gallus* and used that sequence to extract the orthologous region from other galliform genome assemblies using blastn (Camacho et al. 2009).

The sequences from each of these species was aligned using MUSCLE (Edgar 2004b, a) as implemented on the EMBL-EBI web services site (https://www.ebi.ac.uk/Tools/msa/muscle/; see Madeira et al. 2019) using default parameters. We used Mesquite v. 3.6.1 (Maddison and Maddison 2021) to manually identify exon boundaries using the annotated chicken genome. We removed a section at the 3’ end of *LYZ* of *Penelope pileata* because it appeared highly divergent. Then we removed all gap columns (created by exclusion of taxa) and corrected misalignments (to match exon-intron boundaries) manually. We identified three long autapomorphic insertions in intron 2 (in *Callipepla squamata, Penelope pileata*, and *Odontophorus gujanensis*). These long insertions were excluded from analyses, and the data was re-aligned manually after exclusion of these regions prior to analyses to ensure no nucleotides in other species were aligned with any excluded region.

After these modifications, we created several datasets for analyses: 1) the entire region; 2) the pre-*LYZ* region (the section upstream of the coding region); 3) the combined coding exons (introns excluded); 4) intron 1; 5) and intron 2. This was done to separate out different regions that may show different evolutionary histories, or patterns due to variation in mutation and selection rates. Intron 3 is relatively short (332 bp) and so was not analyzed as an individual partition.

### LYZ tree estimation

The maximum likelihood (ML) tree for each of our *LYZ* datasets (total, as well as the individual components) was estimated in IQ-TREE using the “TEST” option, with 1000 ultrafast bootstrap replicates (Hoang et al. 2018) to assess support.

### Identifying sites that support alternative topologies

We determined which nucleotides supported one of two alternative topologies as a way to identify sites (or regions) that might support different sets of relationships. The focus was to compare an alternative tree which placed New World quail and guineafowl as a monophyletic group (indicated a single TIS→SVT transition), or the likely species tree where these groups were not united (e.g., Fig. 1b). Since our estimated species tree differed from the estimated lysozyme tree in one clade within Phasianidae (see results), we modified the species tree to match the lysozyme tree for that clade so the only difference was in the relationships of interest for understanding lysozyme evolution. We compared the site likelihoods for each site given these two different topologies. Site likelihoods were obtained using IQ-TREE implementing the *-wsl* command. For each site in the alignment, we subtracted the likelihood value of the alternative topology from the species tree topology so that positive values were sites that favored the species tree and negative values were those that reflected the alternative topology. For this data, we considered all sites, but also identified “decisive” sites as those where the difference in lnL was greater than 5 standard deviations (Kimball et al. 2013).

### Testing for recombination

To determine whether we could find signatures of recombination between the three amino acids of interest, we used GARD (Kosakovsky Pond et al. 2006a, b) as implemented on the DataMonkey server (Weaver et al. 2018). We used the entire alignment (excluding the autapomorphic insertions) with default parameters. Using the *c*-genes (recombination-free segments with a specific gene tree) identified by GARD, we also estimated gene trees using the same methods as described above for the different *LYZ* datasets.

### Data availability

Lysozyme alignments and treefiles generated are included in supplementary material.

## Results

### The LYZ locus tree conflicted with the species tree

An estimate of the Galliformes species tree (Fig. 2a) generated using UCE data scattered throughout the genome had a topology consistent with expectation based on published multigene and phylogenomic trees (Fig. 1c). The coalescent branch separating guineafowl from the Odontophoridae + Phasianidae clade was short (0.63) (Fig. 2a). In contrast, the coalescent branch lengths that unite most galliform families were longer (>4). This favors ILS as an explanation. In agreement with this, the distribution of the SVT sequence in lysozymes could be reconciled with this species tree if the topology of the *LYZ* gene tree placed pheasants outside of a New World quail + guineafowl clade. Using UCEs, that alternative topology has 18.7% quartet support (Fig. 2a), indicating that many UCE gene trees have this topology. The quartet support for the two minority topologies associated with each internal branch, including the focal branch that unites New World quail and guineafowl exhibited modest asymmetry in almost all cases. This suggests that ILS is likely to represent the primary explanation for discordance among gene trees for the focal node.

**Fig. 2.**
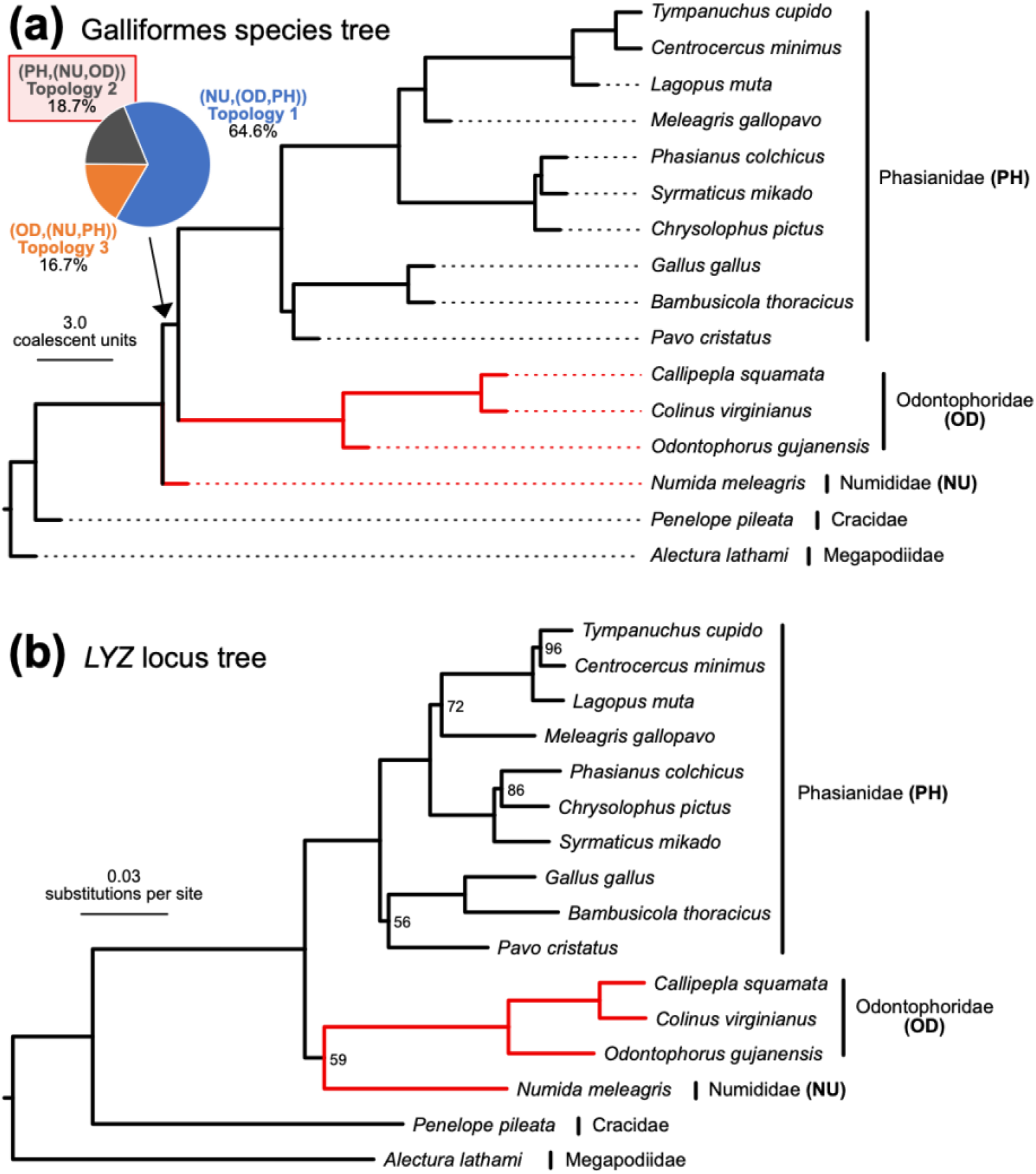
ILS can explain a single gain of the SVT variant. (**a**) Estimate of the Galliformes species tree generated by wASTRAL analysis of 4284 UCE gene trees. Red branches indicate taxa with the SVT variant; other taxa have the TIS variant. All internal branches had maximal (1.0) local posterior probabilities. The pie chart presents quartet frequencies for the three possible topologies for the focal branch. We refer to topology of the wASTRAL tree as topology 1 and the two conflicting topologies as 2 and 3. Topology 2 is highlighted in red because it unites taxa with the SVT variant. Internal branches reflect coalescent units and terminal branch lengths are arbitrary. (**b**) The ML estimate of the *LYZ* locus tree based on analysis of the complete region is topology 2. Ultrafast bootstrap support values <100% are presented to the right of all nodes in the tree; nodes without values have complete (100%) support. Treefiles and quartet support for all branches are available as Supplementary Files 2 and 3, respectively.

The ML estimate of the phylogeny for the entire *LYZ* gene region (Fig. 2b) was consistent with a single origin of the SVT variant and, therefore, conflicted with the species tree. While this topology only required a single TIS→SVT transition, support for the Numididae + Odontophoridae clade was low (61%) in the *LYZ* locus tree given the length of the total *LYZ* alignment (Supplementary Table 1). One explanation for this limited support would be intralocus recombination, a hypothesis that predicts that subsets of the *LYZ* gene region might support alternative trees. This is the reason we referred to the tree in Fig. 2b as a locus tree; the number of gene trees underlying this region is an open question.

Dividing the *LYZ* alignment into functionally-defined subsets corroborated the hypothesis that there were likely multiple gene trees underlying the *LYZ* locus (Supplementary Table 1). Alignments comprising the exons and introns in the middle of the locus supported a tree that united Numididae with the Odontophoridae (i.e., agreed with the overall locus tree) but analyses of ∼2000 bp upstream of the start codon supported a tree congruent with the Galliformes species tree (Supplementary Table 1). These results suggest that there was intralocus recombination, with at least part of the *LYZ* coding region having the species tree topology for the focal node (approximately 64.6% of UCE gene trees; Fig. 2a) and another part having the alternative tree that unites Numididae and Odontophoridae (consistent with 18.7% of UCE gene trees; Fig. 2a).

### Varying site patterns and recombination breakpoints in the LYZ locus

To further explore topological conflicts within the *LYZ* gene region, we examined differences in the site likelihoods given the species tree and the optimal tree for the *LYZ* gene region (Fig. 3a). Site likelihoods can identify specific locations, or regions, that may favor one topology over another, with decisive sites (Kimball et al. 2013) showing sites that strongly favor one topology. The upstream region and the end of the locus (intron 3 and exon 4) on average favored the species tree (Fig. 3a and Supplementary Table S3), while the other regions (exon 1 through exon 3), on average, supported the alternative topology. We identified 46 sites as being decisive; these were scattered throughout the entire locus (except exon 4) and followed the same pattern as sites overall (Supplementary Table S3). Two of the decisive sites that supported the *LYZ* locus tree corresponded to first codon positions of the focal amino acids in exon 2 (amino acids 40 and 56) that unite Numididae and Odontophoridae; the codon for the third focal amino acid had little support for either the species phylogeny or the *LYZ* gene tree (Fig. 3a and Supplementary Table S4). The limited support for either tree in the codon for amino acid position 91 (which specifies either Ser or Thr, depending on the galliform family) is likely to reflect the fact that the ancestral state is a TCN serine codon and the derived state within Phasianidae is an AGY serine codon.

**Fig. 3.**
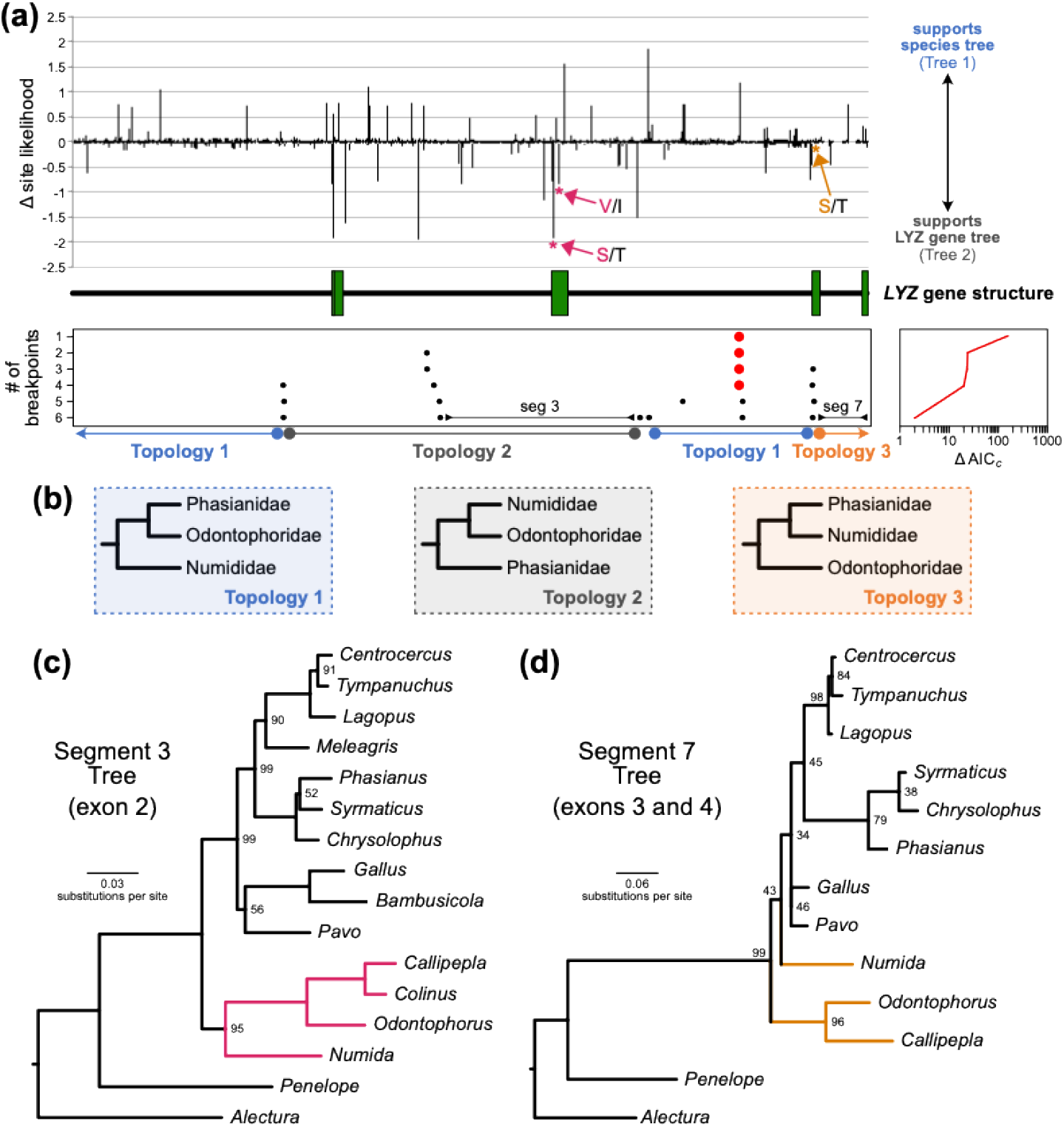
Evidence for recombination during the evolution of the LYZ gene region. (a) Map of the LYZ gene (exons indicated using green boxes) with differences in site likelihoods given two candidate trees (the species tree and the LYZ locus tree) shown above the locus map and potential recombination breakpoints based on GARD analysis are shown below the map. The positions of the codons for the focal amino acids are indicated on the site likelihood graph. Differences in model fit (AICc) given each breakpoint number is presented to the right of the breakpoint graph. The topology for the focal node is presented below the recombination breakpoint map and the segments containing the exons with the focal amino acids are labeled “seg 3” and “seg 7.” The topology and segment maps assume six breakpoints (the best-fitting model in GARD). (b) The three relevant topologies for relationships among guineafowl, New World quail and pheasants. Colors match the topology map in part a. (c) Gene trees estimated using alignment segment 3 and alignment segment 7, indicating the exons located in each segment. Branch colors indicate the state for the focal amino acids (i.e., S and V for segment 3 and T for segment 7). Ultrafast bootstrap support is indicated (unless it is 100%)

Since the decisive sites favoring either of the candidate trees were somewhat intermixed, the likely positions of any recombination breakpoints were unclear based on this analysis. To examine the possibility of intralocus recombination in greater detail, we conducted an analysis using GARD, which identifies potential recombination breakpoints and divides alignments into segments with different phylogenies (*c*-genes). GARD examined over 95,000 models with between one to six breakpoints (Fig. 3a). The model considering at least one recombination event was substantially better than assuming no recombination (ΔAICc = 248.27). The best model had six breakpoints (AICc = 58236.95), though five breakpoints were also credible (ΔAICc = 1.99). Models with fewer breakpoints were all much worse (ΔAICc >8). The models with five versus six breakpoints were very similar (breakpoints were within a few base pairs of each other), with the exception that including six breakpoints added a third breakpoint within intron 2, creating a small (72 base pair) *c*-gene.

Regardless of how many breakpoints were considered, all models (even the single breakpoint) suggest recombination within intron 2, which is the intron between exon 2 (which encodes the first two focal amino acids) and exon 3 (which encodes the third focal amino acid). The breakpoint with the strongest support was located in intron 2 (red dots in the Fig. 3a breakpoint graph), though the exact position of the breakpoint shifted slightly in models with five versus six breakpoints.

When estimating gene trees for the different regions (*c*-genes) defined by the estimated recombination events (Table 1), different topologies were obtained (though some of the topological differences observed for trees estimated using different alignment segments, particularly shorter segments, suggest some gene tree estimation error). We note that all three of the possible topologies for the families in Phasianoidea (Numididae, Odontophoridae and Phasianidae), were recovered when the segments defined by the six-breakpoint model were analyzed (Fig. 3b). Exon 2 is in a region (segment 3) with the same topology as the overall *LYZ* locus tree whereas exon 3 is in a region (segment 7) with topology 3 (Fig. 3c).

**Table 1.** Topologies and bootstrap support for relevant node *c*-genes defined by GARD. Results presented for the best model, with six breakpoints (results using four and five breakpoints can be found in Supplementary Table S5). Support provided for the branch uniting the two derived families for each topology (see Fig. 3b).

| Dataset | Aligned Length (bp) | Topology 1 | Topology 2 | Topology 3 | Regions included |
| --- | --- | --- | --- | --- | --- |
| Segment_1 | 1753 | 78% | -- | -- | 5' Upstream |
| Segment_2 | 1345 | -- | 64% | -- | 5'+ Ex1+ Int1 |
| Segment_3 | 1662 | -- | 95% | -- | Int1 + Ex2 + Int2 |
| Segment_4 | 72 <sup>a</sup> | -- | -- | -- | Int2 |
| Segment_5 | 792 | 82% | -- | -- | Int2 |
| Segment_6 | 598 | 81% | -- | -- | Int2 |
| Segment_7 | 487 | -- | -- | 43% | Ex3 + Int3 + Ex4 |

<sup>a</sup> No tree estimated due to segment length

## Discussion

Our results provide a resolution to the long-standing enigma surrounding lysozyme evolution. While the initial observation of a conflict between the organismal phylogeny and lysozyme sequences has been explained by convergence or parallelism (Ibrahimi et al. 1979), the Malcolm et al. (Malcolm et al. 1990) experiments suggested it was unlikely the *LYZ* gene underwent two sets of transitions between the TIS and SVT variants. Our results confirmed earlier hypotheses that the species tree differed from the LYZ gene tree. However, our species tree also indicated that a relatively large number of polymorphisms were retained over the branch uniting Odontophoridae and Phasianidae to the exclusion of Numididae (i.e., that there is substantial conflict between gene trees and the species tree due to ILS for this branch).

Additionally, we noted that the trees estimated from separate functional regions of *LYZ* and, more importantly, from distinct *c*-genes (defined by estimated recombination break points) revealed different evolutionary histories within the *LYZ* gene. This topological conflict among the subregions was not driven exclusively by the sites associated with the three relevant amino acids. If the codons for the three functionally-relevant amino acids had been the strongest decisive sites it would have suggested a phylogenetic artifact driven by convergence; the observation that many other decisive sites supporting different topologies were identified support genuine conflict among gene trees. Our results also provide strong evidence that recombination occurred between the two relevant exons (exons 2 and 3) occurring during the time when the polymorphism was present.

### A model of LYZ evolution in Galliformes reflecting ILS and recombination

Although our results indicate that ILS is an important part of the explanation for the conflict between the distribution of lysozyme types and the galliform species tree, discordance due to ILS is only part of the explanation. Indeed, the simplest way to explain the distribution of two lysozyme types using ILS is almost as puzzling as the scenario implicit in Malcolm et al. (1990). ILS requires the maintenance of polymorphisms over multiple speciation events; it is unlikely that deleterious alleles will be maintained in a polymorphic state for long periods of time. The resolution to the problem reflects recombination, leading to the existence of multiple *c*-genes (genomic segments with a single phylogenetic history; Doyle 1997) in the *LYZ* gene region. We found evidence for all three of the possible topologies for phasianoid families (i.e., Numididae, Odontophoridae and Phasianidae) represented in different parts of the *LYZ* locus. Exon 2 (which encodes Thr/Ser 40 and Ile/Val 55) and exon 3 (which encodes Ser/Thr 91) are located in distinct *c*-genes (Fig. 3c,d) that have topologies 2 and 3 respectively (neither of which correspond to the species tree, which we have designated topology 1). This indicates that initial mutant alleles in those exons likely arose independently and underwent recombination to generate the SVT variant of Numididae and Odontophoridae (i.e., allowing lysozyme to move between TIS and SVT more quickly than expected given successive mutations in the same background).

Our ILS + recombination model for galliform *LYZ* evolution (Fig. 4) makes it possible to infer more information about the ancestral states than the simple mutation model implicit in Malcolm et al. (1990). The order in which specific amino acid changes occurred is completely unclear in the Malcolm et al. (1990) scenario. In contrast, the recombination model implies that the initial exon 2 and exon 3 mutations arose independently (Fig. 4). This implies that a TIT lysozyme was present in an ancestral galliform population because the only focal amino acid in the *c*-gene we designate segment 7 (Fig. 3d) is Ser/Thr-91. The most parsimonious evolutionary pathway for amino acid 91 given the local gene tree involves the ancestral serine, encoded by a TCN codon, that undergoes a non-synonymous T□A first position mutation (leading to the threonine present in Numididae and Odontophroidae. Then threonine underwent a C□G second position mutation generating the AGY-encoded serine in Phasianidae (Supplementary Fig. S2).

**Fig. 4.**
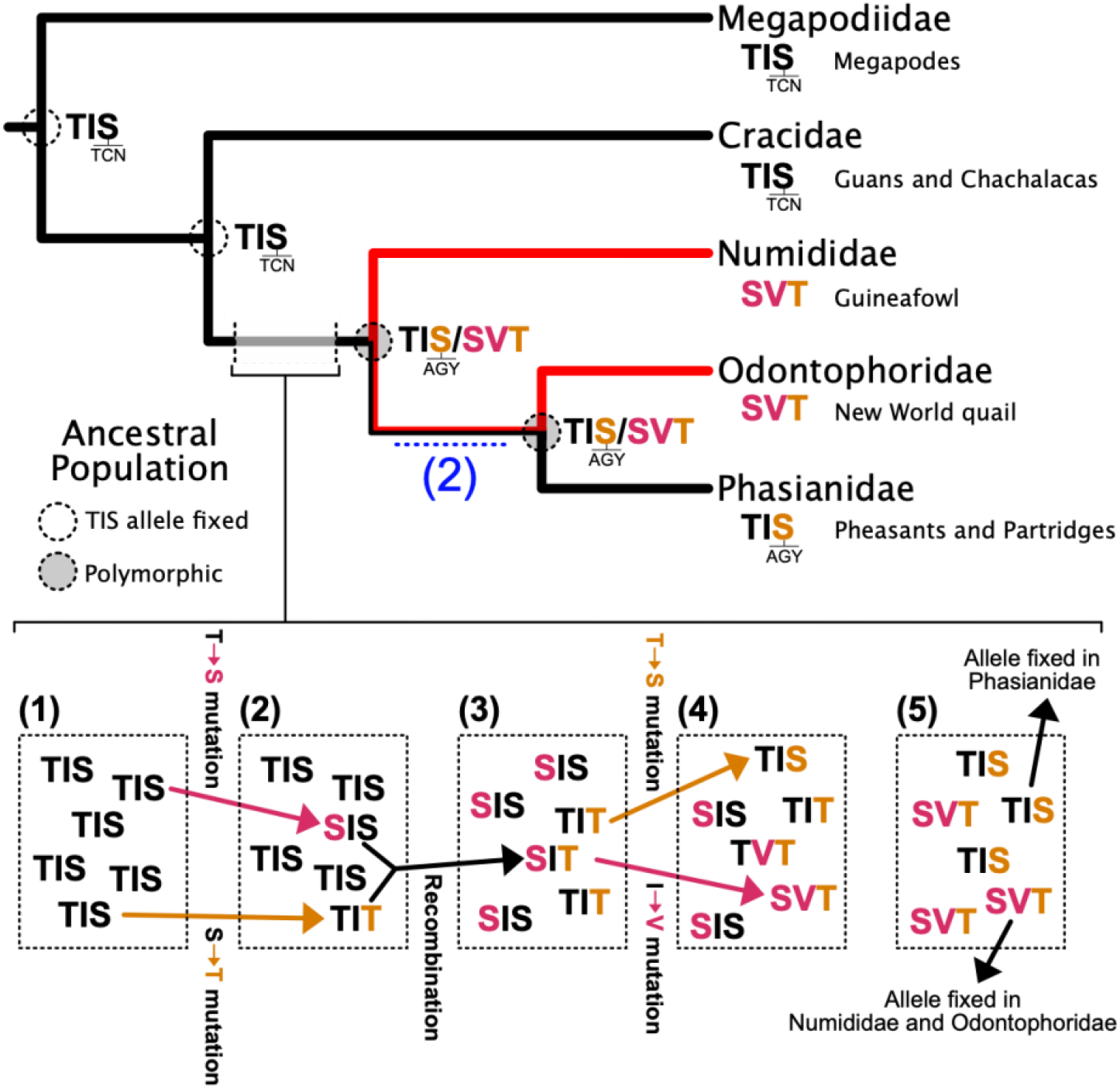
Model for evolutionary changes in the lysozyme gene present in ancestral galliform populations. States for internal nodes reflect populations (node circles) inferred using the polymorphism parsimony criterion, while the blue “2” denotes Scenario 2 in Figure 1. This criterion suggests that all relevant changes occurred on the branch uniting Phasianoidea (Numididae, Odontophoridae, and Phasianidae). We indicate the codons for the serine residue in exon 3 (TCN or AGY). We illustrate five steps: 1) we begin with the ancestral population, with the TIS_(TCN)_ variant fixed; 2) independent mutations generate segregating alleles of TVS_(TCN)_ and TIT; 3) recombination generates a TVT allele; 4) independent mutations generate the SVT allele from an individual with the TVT allele and the TIS_(AGT)_ allele from an individual with the TIT allele; 5) the frequency of the TIS_(AGT)_ and SVT allele increase and they dominate the ancestral population. The common ancestors of all Phasianoidea and of Odontophoridae+Phasiandae were state 5 and state 5 must be maintained over the branch uniting Odontophoridae+Phasiandae. The postulated history indicates the ordering of events rather than the precise timing

The first mutation to occur in exon 2 is uncertain; the TVS variant exhibits the lowest *T*_*m*_ and *T*_*i*_ (presumably the worst thermal properties of any single mutant but the SIS single mutant exhibits the lowest enzymatic activity (Supplementary Fig. S1), Therefore, the amino acid most likely to be present in the in the first exon 2 mutant depends on the biochemical parameter that best correlates with fitness (assuming the novel allele is neutral or only slightly deleterious).

However, the SVT allele arose by recombination and a second non-synonymous mutation in exon 2. The order of the recombination and second exon 2 mutation also cannot be established with absolute confidence, although the SVS variant (the exon 2 double mutant) has poor enzymatic activity and thermal properties (Supplementary Fig. S1) making early recombination between either a TVS or SIS allele and a TIT allele the more likely scenario (the TVS/TIT recombination pathway is shown in Fig. 4 as an example).

There are two major challenges for understanding the patterns of lysozyme evolution given the results of Malcolm et al. (1990). First, the biochemical properties of lysozyme that are responsible for selection in natural populations are unclear. Second, substitutions at other sites have the potential to alter the biochemical phenotype expected given the three focal amino acids. It is clear that selection cannot involve *T*_*m*_ or *T*_*i*_ directly because those parameters only become relevant at temperatures in excess of 50°C. However, other aspects of lysozyme biochemistry are likely to correlate with *T*_*m*_ or *T*_*i*_ and those other parameters may be subject to selection. Shih and Kirsch (Shih and Kirsch 1995) found a number of additional residues that have an impact on both the thermal and enzymatic properties of chicken lysozyme and changes in those residues may compensate for changes in the amino acids identified by Malcolm et al. (1990). Indeed, compensatory mutations may explain variation in the focal residues outside of Galliformes.

### Are there plausible alternative models of lysozyme evolution in Galliformes?

There are other possible explanations for the evolution of LYZ than invoking both ILS and recombination, such as we did in our model. However, they are less parsimonious given our observations.

One alternative would be to postulate that the inferences based on tree topologies were incorrect. The simplest version of this alternative hypothesis postulates that the galliform species tree only requires a single transition; this would require the species tree to have topology 2 (Fig. 3b). This hypothesis motivated Kornegay et al. (1993) to estimate a galliform phylogeny using the mitochondrial cytochrome *b* gene. Kornegay et al. (1993) recovered topology 3, which requires two transitions between the variants. Subsequent large-scale (including whole genomes) data collection and analyses show that both the mitochondrial tree (Meiklejohn et al. 2014; Kimball et al. 2021) and the nuclear species tree (Kimball and Braun 2014; Hosner et al. 2016, 2017; Stiller et al. 2024) for galliforms have topology 1. However, topologies 1 and 3 both require two TIS→SVT transitions (consider the distribution of SVT variants relative to the topologies in Fig. 3b). Regardless, the strong support from topology 1 as the species tree from this and other studies make it unlikely that the Galliformes species tree is incorrect.

Another alternative would be to postulate that the transition between the TIS and SVT variants was crossed very rapidly, such that there was little time for selection to remove intermediate, deleterious alleles. This could occur either due to a “composite mutation” or a set of mutations that occurred in rapid succession. Composite mutations, defined as cases where multiple *de novo* mutations occur in relatively close proximity, do occur (Schrider et al. 2011; Besenbacher et al. 2016) but they represent ∼3% of *de novo* mutations. Thus, it is highly unlikely that such an event would require not just two, but three non-synonymous changes that affect functionally important residues occurring rapidly, particularly as (unless ILS is invoked), this unlikely scenario would have to occur twice.

Finally, it is possible that the conclusion by Malcolm et al. (1990) that intermediates between TIS and SVT are outside of the thermal neutral zone and are selected against is simply incorrect. Malcolm et al. (1990) defined the thermal neutral zone based on the thermal properties of TIS and SVT. Later work looking directly at enzyme activity suggested that TVS, SIT, and TVT all fit within the neutral zone defined by activity levels (Shih et al. 1995). Using this, going from TIS □TVS □TVT □ SVT (or the reverse) could be accomplished using steps of single substitutions while remaining in the enzyme activity neutral zone. However, considering the number of substitutions required by each pathway, this scenario requires six substitutions (three for TIS to SVT, and three to reverse this, accommodating the different serine codon found in the TIS in Phasianidae; each amino acid substitution requires just a single nucleotide substitution). In contrast, our model requires only four substitutions (again, with a single nucleotide substitution required for each amino acid substitution). Assuming an ILS-only model in which topology 2 represents the *LYZ* gene tree requires five nucleotide substitutions (though only four amino acid substitutions, as two nucleotides substitutions are required to change the TCN serine to an AGY serine). Thus, our model requires fewer substitutions. Additionally, the conclusions of Malcolm et al. (1990) were also based upon the observation that intermediate variants had not been observed in extant taxa. This remains true even as more galliform genomes are sequenced; we were unable to find any sequenced LYZ from galliforms that had any of the possible intermediates (Supplementary File 4).

Our proposed recombination model corroborates the Malcolm et al. (1990) hypothesis that intermediates are deleterious in natural Galliformes populations, at least in the case of the TIT variant. Our model indicates that a TIT allele must have been present in an ancestral population and that allele had two fates: 1) back mutation to TIS (with the serine encoded by an AGY codon); and 2) it underwent recombination, ultimately leading to the SVT variant in Numididae and Odontophoridae. The first pathway is consistent with the Rogozin et al. (2016) hypothesis that selection favoring conservation of serine at specific positions can drive rapid transitions from Ser(TCN)□Thr(ACN)□Ser(AGY) (Rogozin et al. [2016] also emphasize the existence of a Ser[TCN]□Cys[TGY]□Ser[AGY], but that does not appear to be relevant for this study). The second fate (recombination) further emphasizes that the ancestral TIT allele did not persist for a long period of time. A simple way to explain why additional changes occur rapidly after the origin of an allele is to postulate that the allele has lower fitness and that the additional changes are compensatory.

### Implications for ancestral protein reconstruction

“…once the structures of ancestral polypeptide chains are known, it will in the future be possible to synthesize these presumed components of extinct organisms. Thus one will be able to study the physico-chemical properties of these molecules and to make inferences about their functions.” - Pauling and Zuckerkandl (Pauling and Zuckerkandl 1963).

Similar to Malcolm et al. (1990), many studies that seek to do “paleogenetics” (Pauling and Zuckerkandl 1963), or reconstruction of ancestral protein sequences, do not consider processes such as ILS or recombination. The foundational work on ILS (Hudson 1983; Tajima 1983) was before Malcolm et al. (1990), but the publication that was most influential for highlighting the importance of ILS in phylogenetics (Maddison 1997) was later. Additionally, ILS is primarily viewed as a phenomenon of neutral evolution, so it may not be applicable to most studies that focus on understanding functional changes over evolutionary time (Thornton 2004; Harms and Thornton 2010; Selberg et al. 2021) where selection acts to quickly remove intermediates of lower fitness. In these cases, polymorphisms are unlikely to persist through subsequent speciation events. However, the Malcolm et al. (1990) experiments were ultimately about the paradox of finding that mutational pathways implied by ancestral reconstruction implied deviations from neutral evolution. In this case, the TIS and SVT alleles are likely to have had equal fitness (at least based on the thermal data; Malcolm et al. 1990), so a TIS/SVT polymorphism could have been maintained through multiple speciation events.

We found that recombination played an important role in resolving the problematic mutational pathway, a finding that might have been predicted by classical Fisher-Muller theory (Fisher 1930; Muller 1932) as long as the distance between the functionally-relevant sites are large enough to permit substantial intralocus recombination that combines the higher fitness variants. Mirarab et al. (2024) suggested that realistic recombination rates and effective population sizes for birds implied that recombination and ILS were relevant at scales between 5 and 5000 bp. Those distances are small enough for intralocus recombination to play an important role in molecular evolution. The results of this study are consistent with the idea that recombination on this scale is important in birds.

This study raises an important question if we look beyond lysozyme: should studies focused on the reconstruction of ancestral proteins consider ILS or recombination when they infer the sequences of ancestors? It is tempting to answer affirmatively any question about whether any process that can play a role in sequence evolution should be considered in molecular evolutionary analyses. Moreover, the fact that these analyses added certainty to the pathway by indicating the existence of an ancestral TIT variant might seem to indicate that including recombination in ancestral state reconstruction is highly desirable. However, ancestral sequence reconstruction has been shown to be robust to model misspecification in many cases (Muñiz-Trejo et al. 2025), suggesting that it is often unnecessary to model “full reality” of the evolutionary process in this type of study. If one is primarily interested in sites subject to selection, fixation of new variants may be sufficiently rapid that ILS is not an issue and so the evolution of many proteins may be able to be explained without considering ILS. While recombination is likely to have occurred in the history of all proteins, it also greatly increases the number of potential ancestral sequences. The number of ancestral sequences in play for galliform lysozymes is small, but most modern studies use much more divergent proteins where there may be substantial uncertainty in the ancestral sequences even if one assumes a single underlying history. Nevertheless, consideration of recombination is likely to be warranted in specific circumstances where unexpected results are encountered.

### Conclusions

Since its initial discovery in 1922 (Fleming 1922), lysozyme has been the focus of much scientific research and it was the first (or a very early) protein used in many genetic and biochemical studies. One of these firsts was its use in “paleobiochemical” experiments by Malcolm et al. (1990). That study led to a paradox by suggesting two transitions across a valley in the fitness landscape. Ultimately, resolving the paradox of how it evolved in galliform birds, highlighted the importance of considering the interaction of ILS and recombination in ancestral state reconstruction. While some proteins may show simple, and easily explained, pathways from ancestral to extant proteins, careful assessment (particularly when simpler models do not appear sufficient to explain extant proteins) can highlight the complex historical processes that led to extant sequences, particularly when simpler processes are insufficient.

## Supporting information

Bird image sources

UCE quartet frequences

LYZ alignment for analysis

LYZ tree files

UCE gene trees

Galliform LYZ coding regions

## Acknowledgements

We thank Brittaney Buchanan, Anna Mohr, Dylan Morris, Shelby Palmer, and Manasee Weerathunga for constructive comments on earlier versions of the manuscript. Elisabeth Smith assisted in extracting UCEs for analyses.

## Funding

This research was facilitated by grants from the US National Science Foundation to RTK and ELB (DEB-1118823 and DEB-1655683).

## Notes

### Competing Interest Statement

The authors have declared no competing interest.

