## Supplementary Tables and Figures for "Recombination and incomplete lineage sorting resolve the enigma of lysozyme evolution"

Supplementary Material

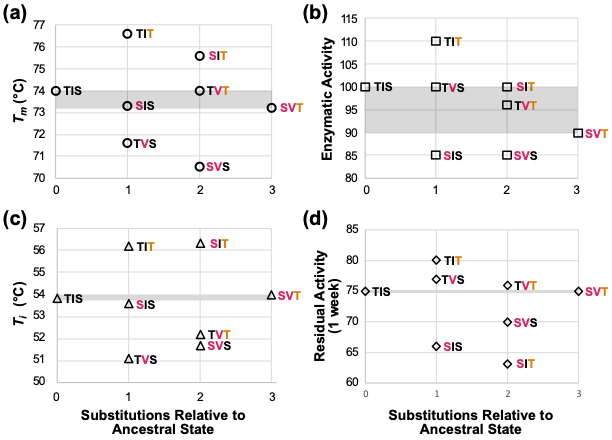

Supplementary Figure S1. Biochemical parameters of extant (TIS and SVT) and ancestral lysozymes.

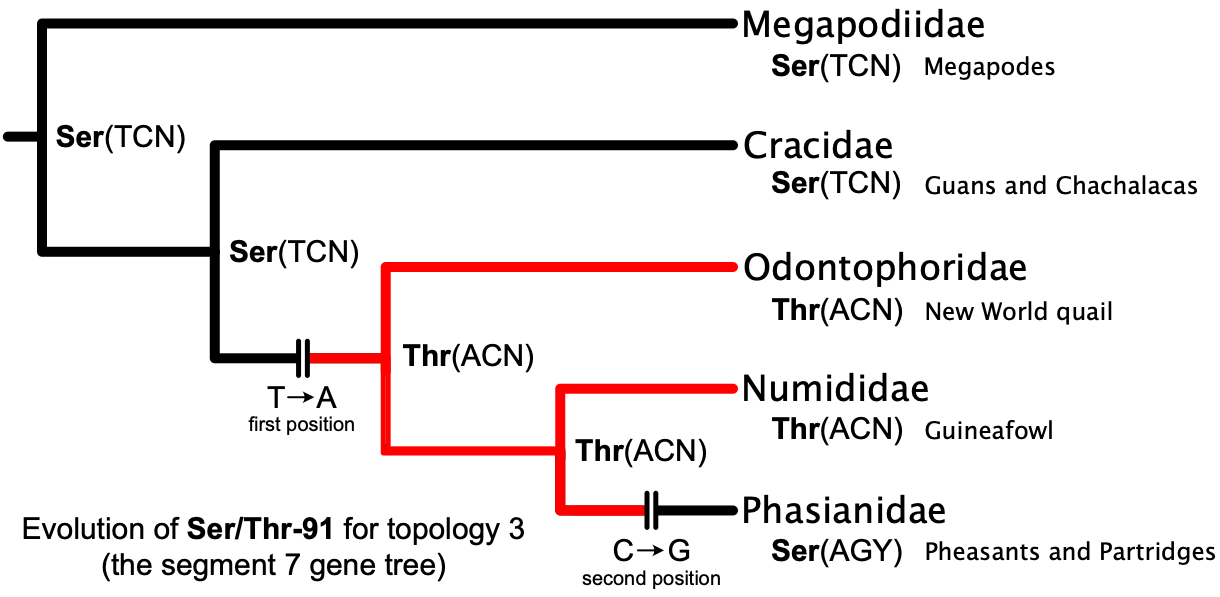

Supplementary Figure S2. Most parsimonious ancestral state reconstruction for Ser/Thr-91 given topology 3. Only non-synonymous substitutions are shown. This pathway is most parsimonious if the ancestral threonine codon is ACY; it is ACC in all Numididae and Odontophoridae that we examined and the serine codons in the Phasianidae we examined vary between AGC and AGY (see Supplementary File 4). We chose topology 3 for ancestral state reconstruction because that is the topology of the *c*-gene that includes the Ser/Thr-91 codon.

Supplementary Table S1. Species used for analysis*.* Names follow AviList (AviList Core Team 2025).

| Family | Species | Common Name |
| --- | --- | --- |
| Megapodiidae | *Alectura lathami* | Australian Brushturkey |
| Cracidae | *Penelope pileata* | White-crested Guan |
| Numididae | *Numida meleagris* | Helmeted Guineafowl |
| Odontophoridae | *Callipepla squamata* | Scaled Quail |
|  | *Colinus virginianus* | Northern Bobwhite |
|  | *Odontophorus gujanensis* | Marbled Wood Quail |
| Phasianidae | *Bambusicola thoracicus* | Chinese Bamboo Partridge |
|  | *Centrocercus minimus* | Gunnison Grouse |
|  | *Chrysolophus pictus* | Golden Pheasant |
|  | *Gallus gallus* | Red Junglefowl |
|  | *Lagopus muta* | Rock Ptarmigan |
|  | *Meleagris gallopavo* | Wild Turkey |
|  | *Pavo cristatus* | Indian Peafowl |
|  | *Phasianus colchicus* | Common Pheasant |
|  | *Syrmticus mikado* | Mikado Pheasant |
|  | *Tympanuchus cupido* | Greater Prairie Chicken |

Supplementary Table S2. Topologies and bootstrap support for the relevant node using *a priori* defined partitions. Support is provided for the branch uniting Odontophoridae and Phasianidae (species topology) or Numididae and Odontophoridae (alternative topology).

| *LYZ* partition | Aligned Length (bp) | Topology1 (e.g., Fig. 2a) | Topology 2 (e.g., Fig. 2b) |
| --- | --- | --- | --- |
| All combined | 6709 | -- | 59% |
| 5' UTR | 2230 | 61% | -- |
| Exons - All | 444 | -- | 81% ^a^ |
| Exons 2 + 3 | 241 | -- | 82% |
| Exons 2, 3, intron 2 | 2296 | -- | 70% |
| Intron 1 | 1648 | -- | 63% |
| Intron 2 | 2055 | -- | 46% |

^a^ Numididae + Odontophoridae nests within Phasianidae

Supplementary Table S3. Summary of site likelihood estimation. Regions that support the species tree are positive and are noted in bold.

| Gene Region | Avg. Delta l*nL* for all sites | Avg. Delta *lnL* for decisive sites |
| --- | --- | --- |
| All combined | -0.0002486 | -0.04553 |
| 5' Upstream | **0.0007228** | **0.12327** |
| Exon 1 | -0.0099546 | -0.43471 |
| Intron 1 | -0.0004184 | -0.01369 |
| Exon 2 | -0.0090426 | -0.31302 |
| Intron2 | -0.0001485 | -0.03024 |
| Exon 3 | -0.0069563 | -0.46240 |
| Intron 3 | **0.0015662** | **0.13886** |
| Exon 4 | **0.0084119** | NA |

Supplementary Table S4. Site likelihoods for key amino acid positions. Decisive sites are in bold. Positive sites support the species tree, while negative sites support the alternative topology.

| Amino Acid | Codon position | Alignment Site | lnL |
| --- | --- | --- | --- |
| 40 | 1 | 4050 | **-1.93713** |
| 40 | 2 | 4051 | 0.00064 |
| 40 | 3 | 4052 | 0.03360 |
| 55 | 1 | 4095 | **-0.86250** |
| 55 | 2 | 4096 | 0.00064 |
| 55 | 3 | 4097 | 0.02790 |
| 91 | 1 | 6258 | -0.04121 |
| 91 | 2 | 6259 | 0.07227 |
| 91 | 3 | 6260 | -0.02946 |

.

Supplementary Table S5. Topologies and bootstrap support for relevant node *c*-genes defined by GARD. Results presented for the best model, with six, five and four breakpoints. Support provided for branch uniting the two derived families for each topology (see Fig. 3b).

| Dataset | Aligned Length (bp) | Topology 1 (see Fig. 3b) | Topology 2 (see Fig. 3b) | Topology 3 (see Fig. 3b) | Regions Included |
| --- | --- | --- | --- | --- | --- |
| **6 Breaks** |  |  |  |  |  |
| Segment_1 | 1753 | 78% | -- | -- | 5’ Upstream |
| Segment_2 | 1345 | -- | 64% | -- | 5’+ Ex1+ Int1 |
| Segment_3 | 1662 | -- | 95% | -- | Int1 + Ex2 + Int2 |
| Segment_4 | 72 ^a^ | -- | -- | -- | Int2 |
| Segment_5 | 792 | 82% | -- | -- | Int2 |
| Segment_6 | 598 | 81% | -- | -- | Int2 |
| Segment_7 | 487 | -- | -- | 43% | Ex3 + Int3 + Ex4 |
| **5 Breaks** |  |  |  |  |  |
| Segment_1 | 1759 | 79% | -- | -- | 5’ Upstream |
| Segment_2 | 1332 | -- | 60% | -- | 5’+ Ex1+ Int1 |
| Segment_3 | 2034 | -- | 80% | -- | Int1 + Ex2 + Int2 |
| Segment_4 | 504 | 100% ^b^ | -- | -- | Int2 |
| Segment_5 | 603 | -- | 70 | -- | Int2 |
| Segment_6 | 477 ^c^ | -- | -- | -- | Ex3 + Int3 + Ex4 |
| **4 Breaks** |  |  |  |  |  |
| Segment_1 | 1752 | 78% | -- | -- | 5’ Upstream |
| Segment_2 | 1296 | -- | 63% | -- | 5’+ Ex1+ Int1 |
| Segment_3 | 2551 | -- | 87% | -- | Int1 + Ex2 + Int2 |
| Segment_4 | 617 | 89% | -- | -- | Int2 |
| Segment_5 | 493 | -- | -- | 44% | Ex3 + Int3 + Ex4 |

^a^ No tree estimated due to segment length

^b^ New World quail+Pheasants are united with guans

^c^ None of these topologies were identified.
