## Supplementary material for "Recombination and incomplete lineage sorting resolve the enigma of lysozyme evolution": Bird image sources

All bird images are 19th/early 20th century paintings available from Wikimedia Commons.

Complete information for each image is listed below:

****

Megapodiidae (Megapodes)

Eulipoa

Eulipoa wallacei

https://commons.wikimedia.org/wiki/Category:Megapodiidae_illustrations?uselang=lt#/media/File:PaintedMegapode_white_background.jpg

Ogilvie-Grant, W. R. (William Robert), 1863-1924 - Handbook to the Game-birds. Volume 2

Viešo naudojimo

File:PaintedMegapode white background.jpg

Sukurta: 1897 m. sausio 1 d.

****

Cracidae (Guans and chachalacas)

Mitu

Ourax mitu » = Mitu mitu (Alagoas Curassow) - adult male

https://upload.wikimedia.org/wikipedia/commons/9/90/Mitu_mitu_white_background.jpg

Nicolas Huet - Nouveau recueil de planches coloriées d'oiseaux

Public Domain

File:Mitu mitu white background.jpg

Created: 1 January 1838

****

Numididae (Guineafowl)

Numida

Numida meleagris » = Numida meleagris (Helmeted guineafowl)

https://upload.wikimedia.org/wikipedia/commons/e/e0/Keulemans_Onze_vogels_1_57_white_background.jpg

John Gerrard Keulemans - Onze vogels in huis en tuin

Public Domain

File:Keulemans Onze vogels 1 57 white background.jpg

Created: 1 January 1869

****

Odontophoridae (New World quail)

Colinus virginianus

https://commons.wikimedia.org/wiki/File:The_Auk_(1898)_(Colinus_virginianus).jpg

Date	1898

Source	The Auk

Author	American Ornithologists' Union

Permission

(Reusing this file)	public-domain: published in the United States pre-1923, also reproduction of art-work where artist died more than 70 years ago.

****

Phasianidae (Pheasants and partridges)

Gallus

Red Junglefowl (Gallus gallus)

https://upload.wikimedia.org/wikipedia/commons/3/38/Red_Junglefowl_by_George_Edward_Lodge_white_background.png

George Edward Lodge - A Monograph of the Pheasants, volume 2 by William Beebe

Public Domain

File:Red Junglefowl by George Edward Lodge white background.png

Created: 1 January 1921
